# Progerin triggers a phenotypic switch in vascular smooth muscle cells that causes replication stress and an aging-associated secretory signature

**DOI:** 10.1101/2022.02.05.479232

**Authors:** Nuria Coll-Bonfill, Urvashi Mahajan, Chien-Jung Lin, Robert P. Mecham, Susana Gonzalo

## Abstract

Hutchinson Gilford Progeria Syndrome is a premature aging disease caused by *LMNA* gene mutation and the production of a truncated lamin A protein “progerin” that elicits cellular and organismal toxicity. Progerin accumulates in the vasculature, being especially toxic for vascular smooth muscle cells (VSMC). Patients’ autopsies show that vessel stiffening, and aortic atherosclerosis is accompanied by VSMC depletion in the medial layer, altered extracellular matrix (ECM), and thickening of the adventitial layer. Mechanisms whereby progerin causes massive VSMC loss and vessel alterations remain poorly understood. Mature VSMC retain phenotypic plasticity and can switch to a synthetic/proliferative phenotype. Here we show that progerin expression in human and mouse VSMC causes a switch towards the synthetic/proliferative phenotype. This switch elicits some level of replication stress in normal cells, which is exacerbated in the presence of progerin, leading to telomere fragility, genomic instability, and ultimately VSMC death. Importantly, calcitriol prevents replication stress, telomere fragility, and genomic instability, reducing VSMC death. In addition, RNAseq analysis shows induction of a profibrotic and proinflammatory aging-associated secretory phenotype upon progerin expression in human primary VSMC. Our data suggest that phenotypic switch-induced replication stress might be an underlying cause of VSMC loss in progeria, which together with loss of contractile features and gain of profibrotic and proinflammatory signatures contribute to vascular stiffness in HGPS. Preventing the phenotypic switch-induced replication stress with compounds such as calcitriol might ameliorate CVD in HGPS patients.

## INTRODUCTION

A-type lamins (lamin A/C) are nuclear structural components that provide a scaffold for the compartmentalization of genome function, and they are involved in the maintenance of genome integrity (van Steensel & Belmont, 2017). Mutations in the *LMNA* gene are associated with degenerative disorders called laminopathies, including Hutchinson-Gilford progeria syndrome (HGPS). HGPS is caused by a heterozygous *de novo* point mutation in the *LMNA* gene (G608G) which results in the synthesis of a truncated prelamin A variant called progerin. Progerin induces multiple cellular and physiological anomalies resulting in growth retardation, loss of subcutaneous fat, alopecia, joint abnormalities, aged-looking skin, and severe cardiovascular disease (CVD) which leads HGPS patients to an early death usually in their teenage years (Filgueiras-Rama et al., 2018; Gonzalo et al., 2017; Hennekam, 2006; McClintock et al., 2007).

Vascular abnormalities in HGPS include atherosclerosis and Vascular Smooth Muscle Cells (VSMC) loss in the aortic media, suggesting a causal connection between VSMC loss and cardiovascular malfunction (Hamczyk et al., 2018; Nevado et al., 2020) Interestingly, the severity of atherosclerosis and VSMC loss correlates with a concomitant increase of Extracellular Matrix (ECM) deposition, leading to vessel stiffening and CVD (Hamczyk et al., 2018). A recent publication demonstrated with a genetic approach, that restoring lamin A ubiquitously in VSMC, ameliorates CVD, indicating that the progerin-induced alteration of VSMC biology is a key contributor to vascular disease in HGPS (Sánchez-López et al., 2021). Moreover, different pharmacological strategies have proved successful in reducing VSMC loss in HGPS mice and improving vascular disease. These strategies include deletion of the matrix metalloprotease 13 (Mmp13) (Pitrez et al., 2020), and the blockade of interleukin-6 signaling in *Lmna*^*G609G/G609G*^ mice (Squarzoni et al., 2021), suggesting that HGPS-associated VSMC depletion involves ECM remodeling and inflammation. Another study has also shown how the disruption of the LINC complex, a protein complex linking the nucleoskeleton and the cytoskeleton, ameliorated aortic disease in progeria mice (Kim et al., 2018). This study highlights that the increased sensitivity to mechanical stress of progerin-VSMC can promote cell death. VSMC loss in *Lmna*^*G609G/G609G*^ mice was also prevented by remodelin, an N-acetyltransferase 10 (NAT10) inhibitor (Balmus et al., 2018), and by an antisense oligonucleotide therapeutic approach using morpholinos targeting progerin expression, which reduced aortic adventitial thickening and extended lifespan (Erdos et al., 2021). However, neither of these studies have fully addressed the reasons behind VSMC loss associated with CVD in HGPS.

Medial mature VSMC from the vascular wall are characterized by low rates of proliferation and migration and their main function is to perform contractile functions, which require proteins such as smooth muscle α-actin (α-SMA), SM22α, calponin, and smooth muscle myosin heavy-chain 11 (MYH11). These cells can undergo a phenotypic switch towards a synthetic phenotype due to different stimuli such as inflammation, oxidative stress, or cell-to-cell contact. This switch is characterized by the downregulation of contractile markers and increased proliferation, migration, and secretion of ECM proteins (Kawai-Kowase & Owens, 2007; Owens et al., 2004; Yap et al., 2021). The thoracic aorta of ≈14-week-old *Lmna*^*G609G/G609G*^ mice shows reduced numbers of α-SMA (measured by hematoxylin staining) and increased deposition of medial collagen that might contribute to cardiovascular alterations in progeria (Del Campo et al., 2019).

At the cellular level, unresolved or unrepaired DNA damage and genomic instability are potent inducers of cell death. Extensive evidence links progerin expression with DNA damage, DNA repair deficiencies, and telomere and mitochondrial dysfunction in different cellular models, including vascular cells (Aguado et al., 2019; Cao et al., 2011; Chojnowski et al., 2015; Coll-Bonfill et al., 2020; Kreienkamp et al., 2018; Kychygina et al., 2021). It is likely that genomic instability contributes to the manifestation of the drastic VSMC loss and CVD in HGPS. For instance, mitochondrial dysfunction and telomeric damage promote VSMC senescence, vessel inflammation, and stiffness (Canugovi et al., 2019; Kychygina et al., 2021; Y. Liu et al., 2013; Y.-F. Liu et al., 2020). However, the molecular mechanisms involved remain unclear. Some studies linked deficiencies in DNA repair involving PARP1 and Endoplasmic Reticulum (ER) stress to cell death in progeria VSMC (Hamczyk et al., 2018; Kinoshita et al., 2017a; Zhang et al., 2014). Replication is one of the major sources of DNA damage that could lead to cell death in dividing cells. We have postulated that persistent genomic instability in VSMC may have dramatic consequences promoting progerin-induced vascular disease.

Here we show for the first time that progerin expression induces a synthetic/proliferative dedifferentiated phenotype in primary human aortic VSMC, which given the inability to properly replicate in the presence of progerin, causes genomic instability and eventually the loss of VSMC. Interestingly, genomic instability emanates to a large extent from replication stress and telomere fragility/dysfunction. Moreover, progerin-induced synthetic phenotype is accompanied by a secretory inflammatory signature associated with fibrosis and premature aging, which could elicit vascular stiffness in HGPS. In addition, we show that calcitriol (the active form of Vitamin D) rescues replication stress and reduces telomere fragility and cell death, positioning calcitriol as a suitable candidate to ameliorate progerin toxicity in the vasculature.

## RESULTS

### Progerin induces a phenotypic switch in human and mouse vascular smooth muscle cells

The mechanisms by which progerin leads to a massive VSMC loss are poorly understood. We hypothesized that the toxic effect of progerin in VSMC could lead to phenotypic switching, as observed in many vascular pathologies (Chakraborty et al., 2021; Coll-Bonfill et al., 2016; Petsophonsakul et al., 2019). To determine the effects of progerin expression in VSMC phenotypic modulation we have used a well-characterized model of human VSMC differentiation based on TGFβ1signaling (Coll-Bonfill et al., 2016; Guo & Chen, 2012). Human primary aortic VSMC (AoSMC) are stimulated with TGFβ1 (10 ng/ml for 24 hours), a growth factor known to promote VSMC differentiation towards a contractile phenotype. In this model, we have monitored AoSMC differentiation analyzing miR-145 expression, a key regulator of contractile/differentiated phenotype in human VSMC (Cordes et al., 2009) (Figure 1A), together with the expression of different contractile genes such as calponin, myoCD, α-SMA or sm22α (Figure 1A). As expected, miR-145 and differentiation markers (myoCD, calponin, and sm22α) increase in cells that were treated with TGFβ1 (Figure 1A), consistent with the acquisition of the contractile VSMC phenotype.

**Figure 1.**
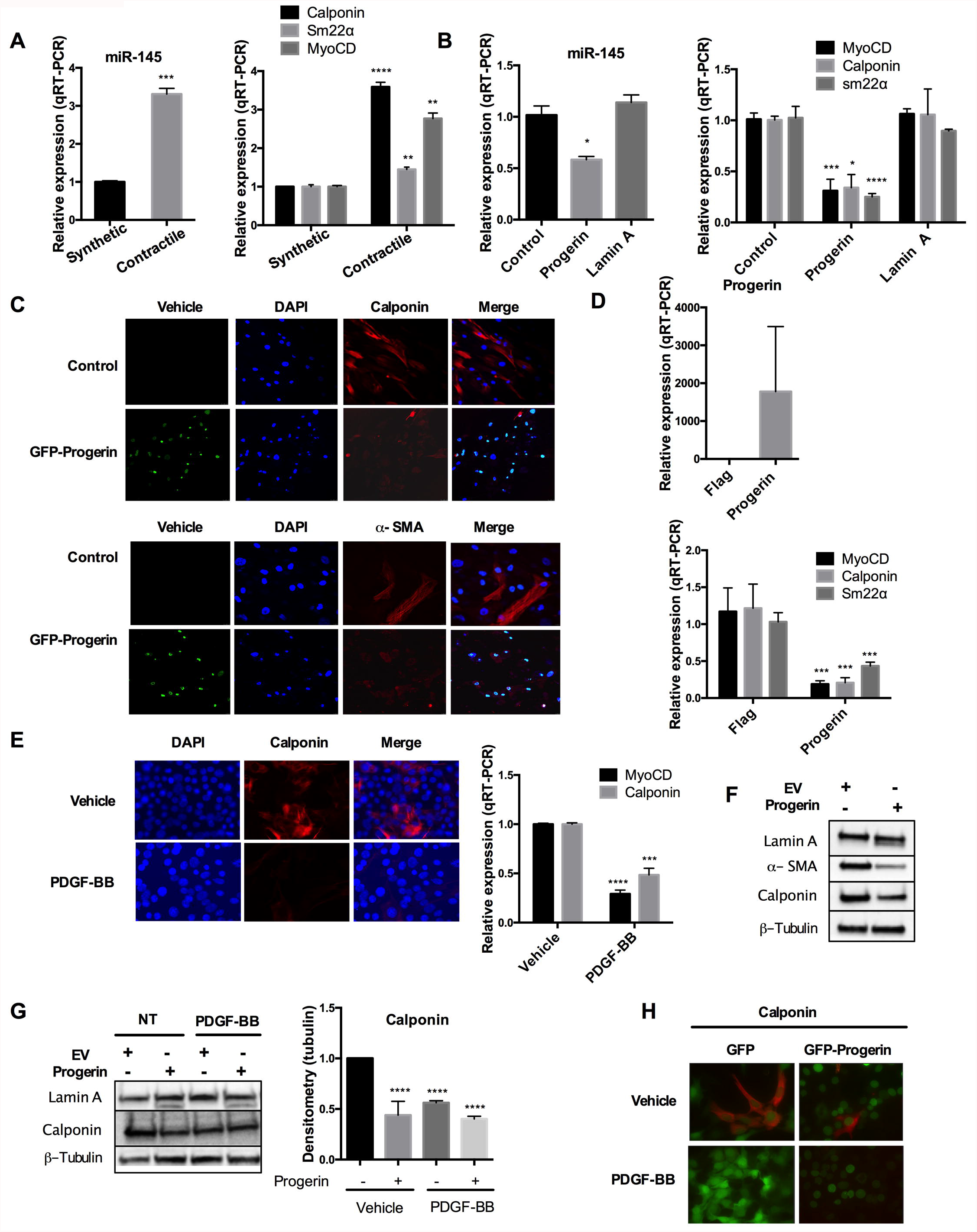
Progerin induces a phenotypic switch in human and mouse vascular smooth muscle cells (MOVAS). A) RT-PCR in Primary Aortic Smooth Muscle Cells (AoSMC) treated with TGFβ1 (10 ng/ml for 24h). Left graph shows levels of miR-145, and right graph expression of contractile proteins. Note the acquisition of contractile/fully differentiated phenotype. Results are average±SEM of two biological replicates in triplicate. B) RT-PCR in AoSMC lentivirally transduced with constructs for doxycycline-induced expression of GFP-progerin/lamin A. Doxycycline (1ug/ml) treatment for 48h showed a decrease in miR-145 expression in GFP-progerin-expressing cells and no changes in GFP-lamin A-expressing cells (left graph). Right graph shows the downregulation of differentiation markers (myoCD, sm22α, and calponin) in GFP-progerin-expressing cells. Results are average±SEM of two biological replicates in triplicate. C) Immunofluorescence of GFP-progerin induction (doxy 1 ug/ml) for 48h showing a reduction of calponin and α-SMA expression. D) RT-PCR in retrovirally transduced AoSMC with Flag or Flag-progerin. Upper graph shows progerin levels and lower graph contractile markers levels (myoCD, calponin, sm22α). Results are average±SEM of two biological replicates in triplicate. E) Representative images of Mouse Vascular Smooth Muscle Cells (MOVAS) treated with PDGF-BB. Note the reduced levels of calponin by immunofluorescence. Right graph shows decreased transcript levels of myoCD and calponin. Results are average±SEM of two biological replicates in triplicates. F) Western Blot in lentivirally transduced MOVAS with pMXIH-progerin and pMXIH-EV (empty vector control). Note the downregulation of α-SMA and calponin upon progerin expression. G) Representative western blot on the left and graph of the densitometry of 3 biological repeats showing decreased levels of calponin on the right. Note that progerin downregulates calponin to the same extent as PDGF-BB. H) Representative images in retrovirally transduced MOVAS with pBABE-GFP and pBABE-GFP-progerin. Note the decrease in calponin levels.

We next evaluated whether progerin can induce an AoSMC phenotypic switch, using a doxycycline (doxy)-inducible GFP-progerin model of AoSMC. The induction of progerin with doxy for 48 hours promoted a significant downregulation of miR-145 (Fig 1B), calponin, sm22 and myoCD transcripts (Figure 1B), indicating a phenotypic switch towards decreased contractility. In contrast, lamin A did not alter the expression of miR-45 or the contractile markers (Figure 1B), indicating that the wild-type protein is not able to induce the phenotypic switch. The reduction of calponin and α-SMA proteins upon expression of GFP-progerin was also evident by immunofluorescence (IF) (Figure 1C). These results were reproduced in AoSMC retrovirally transduced with Flag (control) and Flag-progerin. Progerin-expressing cells presented with a significant downregulation of contractile genes (Calponin, sm22α and myoCD) (Figure 1D). Thus, progerin triggers a phenotypic switch in human primary AoSMC that reduces contractile capabilities.

To determine if the progerin-induced VSMC phenotypic switch is conserved in mouse cells, we used a dedifferentiation model of mouse VSMC (MOVAS) exposed to PDGF-BB, a well-known inducer of dedifferentiation (Holycross et al., 1992; Wang et al., 2004). PDGF-BB induced the downregulation of differentiation markers (myoCD and calponin) by RT-PCR and immunofluorescence (IF) (Figure 1E), as expected. To determine whether progerin also induces a VSMC phenotypic switch in progerin-expressing mouse cells, we lentivirally transduced MOVAS with pMXIH-progerin or pMXIH-EV (empty vector) to obtain a cell line constitutively expressing progerin. MOVAS depicted VSMC phenotypic switch upon progerin expression, showing a decrease in α-SMA and calponin at protein level (Figure 1F). Interestingly, the levels of calponin after de-dedifferentiation using PDGF-BB or progerin expression were very similar, demonstrating how progerin mirrors the capacity of PDGF-BB to promote VSMC phenotypic modulation (Figure 1G).

The same results were observed in a third model in which pBABE-GFP and pBABE-GFP-progerin were retrovirally transduced into AoSMC. Note the reduced positivity for calponin by IF (Figure 1H). Collectively our findings support that progerin induces a switch in VSMC towards a de-differentiated, less contractile phenotype.

### Progerin causes replication stress in VSMC upon the phenotypic switch

In the context of a phenotypic switch induced by progerin, increased proliferation and replication could have detrimental effects, given that progerin causes replication stress in different cellular models (Kreienkamp et al., 2018). Here we tested whether progerin impacts replication of VSMC undergoing de-differentiation and phenotypic switching.

We used MOVAS as a model to study proliferation and DNA replication. PDGF-BB is known to increase proliferation (Gomez & Owens, 2012), while inducing VSMC phenotypic switch in these cells. First, we monitored MOVAS proliferation by ki67 expression upon PDGF-BB exposure, and upon expression of progerin or EV as control (Figure 2A, IF image and graph). Ki67 positivity increased upon PDGF-BB exposure, consistent with increased proliferation. In contrast, proliferation decreased in progerin-expressing cells, as shown in Figure 2A. Progerin also significantly reduced the proliferation induced by PDGF-BB exposure. This result supports the idea that although synthetic VSMC are characterized by high rates of proliferation, progerin expression hinders proliferation in this context. To determine if the reduced proliferation is due to replication stress (RS), one of the main sources of genomic instability that hinders cellular proliferation, we performed single-molecule DNA replication analysis (DNA fiber assays). Replication events were labeled with iododeoxyuridine (IdU) and chlorodeoxyuridine (CldU) for an equal amount of time (20 min). By IF, we measured IdU and CldU tract lengths and calculated the CldU/IdU ratio (Figure 2B). In normal conditions, when there is no RS, CldU and IdU tract lengths are similar and the ratio CldU/IdU is ∼1. However, during RS, events like replication fork stalling/collapse and replication fork instability due to nucleases often result in a shorter second tract (latest replicated DNA) and a ratio CldU/IdU that is less than 1. This is a marker of replication fork instability associated with RS. Interestingly, we find that the average red tract length is higher in progerin-expressing cells than in control cells, indicating an increased speed of the replication fork. In addition, we find a reduced average length of the green tract and a CldU/IdU ratio lower than 1, indicating replication fork instability (Figure 2B). Moreover, analysis of stalled forks (IdU only labeled tracts) showed an increase in progerin-expressing cells (Figure 2C). Our data thus demonstrate that progerin expression in MOVAS causes profound replication stress characterized by fast replication fork speed, fork stalling, and replication fork instability, which hinder proliferation.

**Figure 2.**
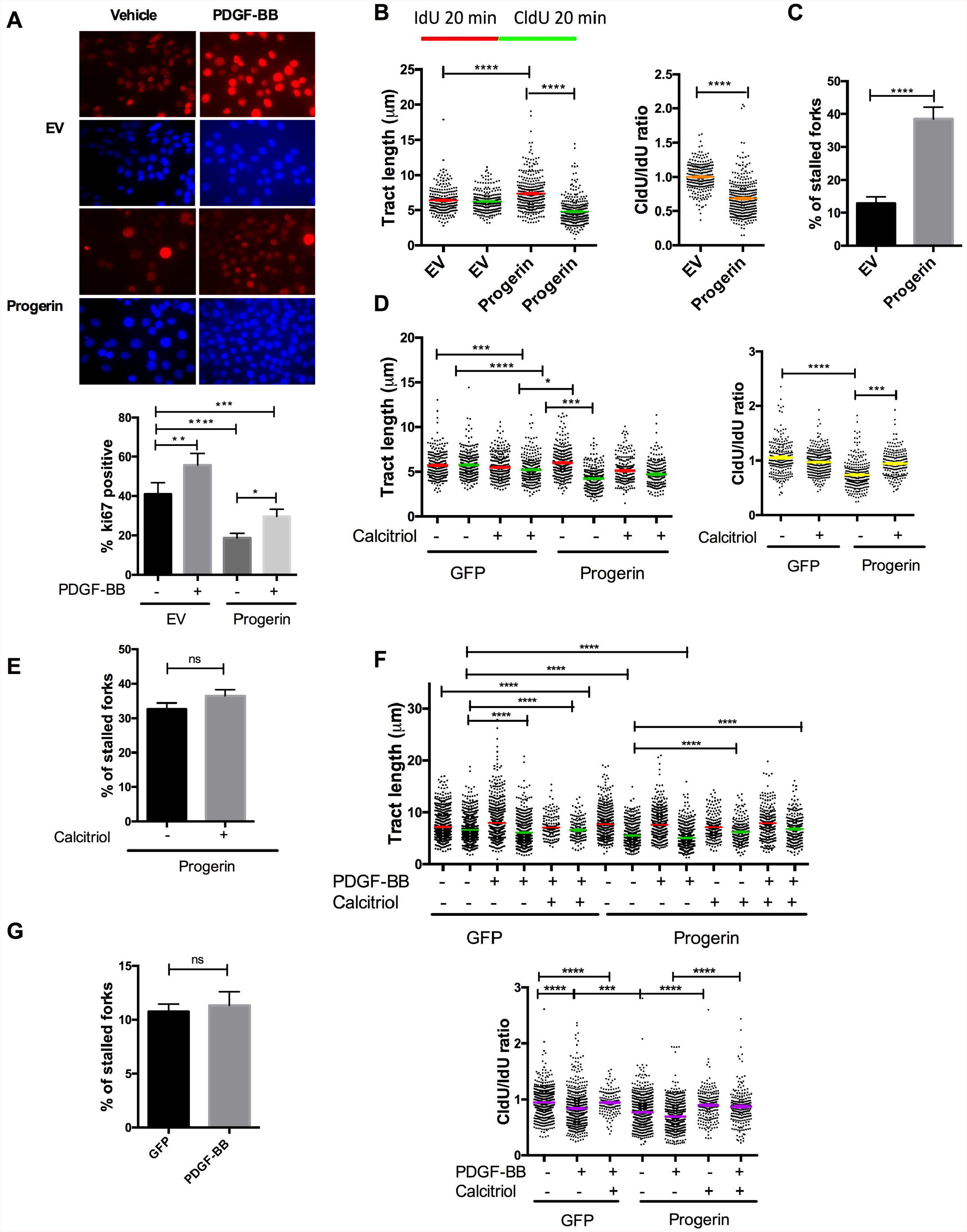
Progerin causes replication stress in VSMC upon the phenotypic switch. A) Representative images of Ki67 immunostaining in MOVAS pMXIH-EV(Empty vector control)/progerin. Right graph shows the quantification of ki67 expression (% positive cells). Results are average ± SEM of three biological replicates. B) Single-molecule DNA replication analysis (DNA Fiber assays) in MOVAS GFP and GFP-progerin cells labeled 20 min IdU and 20 min CldU. Left graph shows tract length, and the right graph shows the ratio between both tract lengths. Note that GFP-progerin cells depict a shortening on the green tract, ratio <1. Results are average ± SEM of three biological replicates. C) Graph shows the quantification of stalled forks (forks that have incorporated only red label from the total of progressing forks). Results are average ± SEM of three biological replicates. D) DNA Fiber assays in MOVAS GFP and GFP-progerin cells +/-calcitriol for 3 days labeled 20 min IdU and 20 min CldU. Left graph shows tract length, and the right graph shows the ratio between both tract lengths. Note that calcitriol rescues tract length in progerin-MOVAS, ratio ≌1. Results are average ± SEM of three biological replicates. E) Graph shows the quantification of stalled forks. Results are average ± SEM of three biological replicates F) DNA fiber assay in MOVAS GFP and GFP-progerin cells +/-calcitriol for 3 days and +/-PDGF-BB, labeled 40 min IdU and 40 min CldU. Left graph shows tract length, and the right graph shows the ratio between both tract lengths. Note that progerin-MOVAS and PGF-BB treated cells depict a shortening on the green tract and a ratio <1 that it is rescued by calcitriol treatment. Results are average ± SEM of four biological replicates. G) The quantification of stalled forks in PDGF-BB. Results are average ± SEM of four biological replicates.

Given that calcitriol (the active form of vitamin D) ameliorates progerin toxicity and RS in other models (Kreienkamp et al., 2018), we tested whether calcitriol could prevent progerin-induced RS in VSMC. MOVAS were treated with calcitriol for 3 days and DNA fiber assays were performed. The results showed that calcitriol rescues RS induced by progerin (Figure 2D, note individual tract lengths and CldU/IdU ratio), but does not reduce fork stalling (Figure 2E). Interestingly, calcitriol was able to slow down replication speed, as shown by a decrease in the red tract length (Figure 2D).

Next, we determined whether RS normally occurs during VSMC phenotypic switch, using MOVAS exposed to PDGF-BB, and how it compares to progerin-induced RS. We show that PDGF-BB increases RS in control cells (Figure 2F, top graph individual tracts and bottom graph CldU/IdU ratio), suggesting that increased proliferation rates promote a certain degree of RS in VSMC. Importantly, PDGF-BB depicted an additive effect in progeria cells (Figure 2F, bottom graph), which suggests a causal connection between increased proliferation and RS. In addition, the red tract is longer in PDGF-BB treated cells and in progerin-expressing cells, indicating an increase in replication fork speed in both cases (Figure 2F). Interestingly, calcitriol slows RF progression (shorter red tracts) and ameliorates RS in PDGF-BB treated MOVAS (Figure 2F) as well. Calcitriol reduces RS in progerin-expressing cells treated or not with PDGF-BB. Moreover, while progerin enhances replication fork stalling, PDGF-BB does not (Figure 2G). Thus, the proliferation of VSMC induced by PDGF-BB causes some RS but this is not as dramatic as the one caused by progerin.

Altogether, the results indicate that progerin hinders replication in VSMC, inducing fork stalling and replication fork instability, which can have dramatic consequences in a cell forced to proliferate due to the phenotypic switch. In contrast, PDGF-BB treated cells show increased proliferation accompanied by relatively modest RS, with no effect in fork stalling, suggesting different mechanisms of replication fork instability induced by progerin and PDGF-BB. Interestingly, calcitriol was able to rescue RS in both cases, seemingly by slowing down replication fork speed.

### Telomeres accumulate replication stress in VSMC undergoing the phenotypic switch

There is robust evidence that progerin expression hinders telomere function, including a decrease in telomere length and increased telomere damage. As testament of the importance of telomere dysfunction driving HGPS phenotype, ectopic expression of telomerase rescues telomere damage, inhibits the DNA damage response at telomeres, improves HGPS cellular phenotypes, and extends lifespan in progeria mice (Aguado et al., 2019; Kychygina et al., 2021; Li et al., 2019).

Telomeres are difficult-to-replicate domains and RS in telomeric regions is one of the principal causes of telomere fragility and dysfunction (Bonnell et al., 2021). To determine whether progerin triggers telomere dysfunction in VSMC during the phenotypic switch, we monitored telomere stability by PNA-FISH staining of metaphase chromosome spreads. Telomere aberrations, such as telomere loss and telomere-containing chromosomal fragments were screened, but no significant difference was observed in progerin-expressing cells compared to control cells. However, fragile telomeres, telomeric regions in which replication is impaired and that are visualized as multi-telomere signals (MTS) were increased in progerin-expressing cells with respect to control cells (Figure 3A, left graph). We observed that in MOVAS, around 50% of the metaphases had 1-4 MTS, indicating basal levels of telomere fragility in these cells. Another 20% of control cells had >5 MTS per metaphase. The remaining 30% of metaphases had no MTS. Expression of progerin increased the amount of MTS per metaphase, so that approximately 40% of the metaphases had MTS over basal levels (>5 MTS per metaphase). Only 10% of metaphases showed no MTS. We also observed increased chromosome end-to-end fusions, primarily Robertsonian Like Fusion (RLF) (Figure 3A, right graph) upon progerin expression; an indication of telomeric dysfunction. These results are consistent with the increase in RS in progerin-expressing cells, linking replication defects and telomere fragility in MOVAS. Then, given that PDGF-BB was able to induce RS, we examined whether PDGF-BB treatment also triggers telomere fragility. However, we find that PDGF-BB does not induce telomere fragility/instability (MTS/RFL). Altogether, the data suggest that MOVAS can maintain telomere homeostasis after PDGF-BB treatment (Figure 3A) but not after progerin expression. Thus, the damaging effect of progerin on replication, telomere stability, and overall genomic stability goes beyond the mechanisms induced by PDGF-BB.

**Figure 3.**
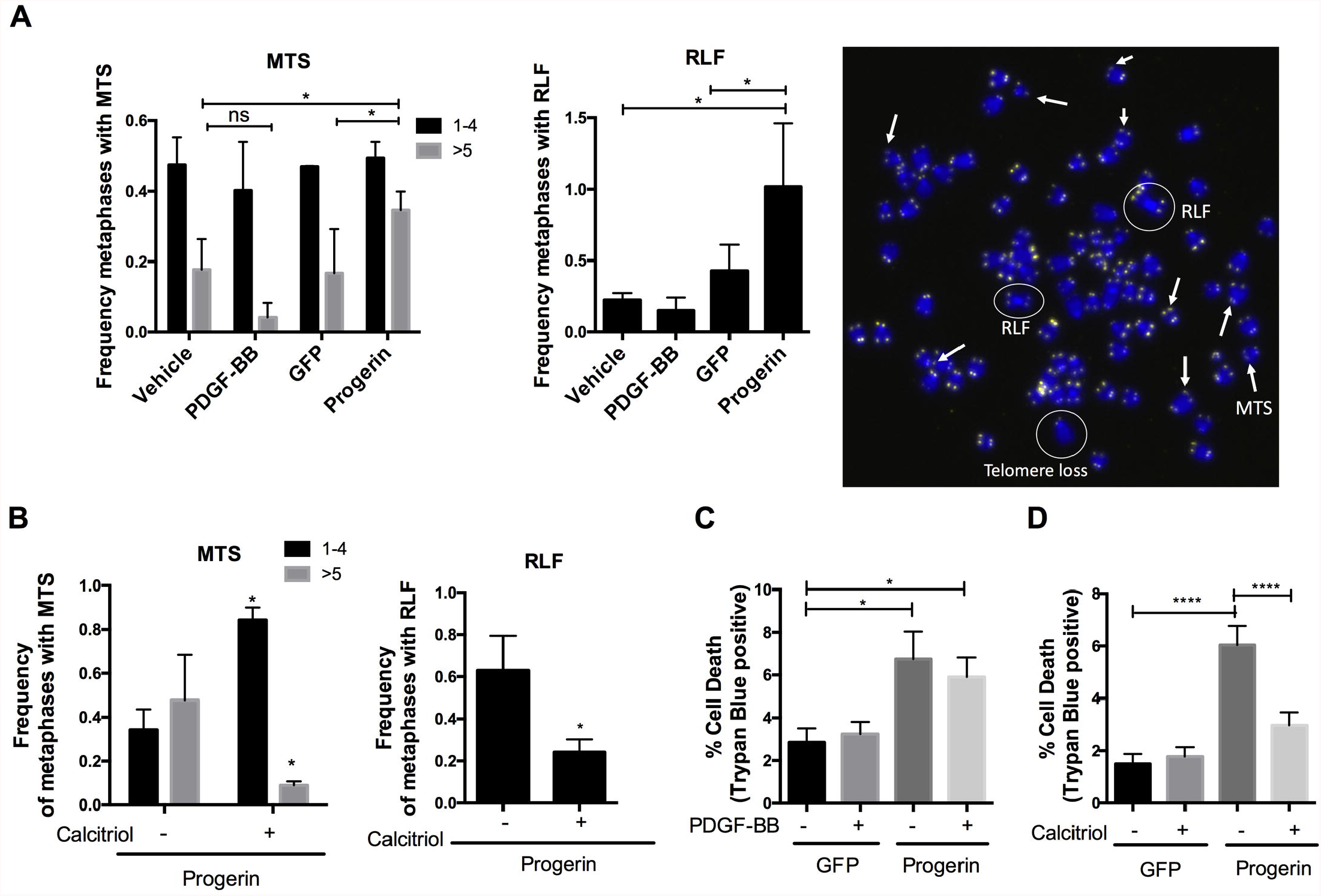
Telomeres accumulate replication stress in VSMC undergoing the phenotypic switch. A) Analysis of metaphase spreads of MOVAS treated with vehicle or PDGF-BB, and MOVAS retrovirally transduced with GFP or GFP-progerin constructs. Graph shows the frequency of metaphases presenting with Multi-Telomeric Signals (MTS) and Chromosomes End-to-End Robertsonian-like fusions (RLF). Around 20 metaphases were analyzed in each experiment. Graphs represent average±SEM of three biological repeats. B) Graphs show frequency of metaphases with MTS and RLF in MOVAS expressing GFP-progerin and growing in culture +/-Calcitriol for 4 weeks. Around 20 metaphases were analyzed in each experiment. Graphs represent average±SEM of three biological repeats. Note that Calcitriol prevented MTS and RLF accumulation during the 4 weeks. C) Graph depicting the quantification of Trypan Blue staining (% of positive cells) on MOVAS expressing GFP or GFP-progerin +/-PDGF-BB. Graph shows the average±SEM of three biological repeats. D) Graph depicting the quantification of Trypan Blue staining (% of positive cells) on MOVAS expressing GFP or GFP progerin +/-Calcitriol. Graph shows the average ± SEM of three biological repeats.

Moreover, since calcitriol rescued RS in progerin-expressing MOVAS, we determined whether calcitriol could prevent the telomere fragility induced by progerin. We treated progerin expressing cells with calcitriol for 4 weeks and monitored the extent of telomere fragility/instability accumulating over time by quantifying MTS and RLF. Progeria cells increased the levels of MTS over time, with approximately 50% of metaphases showing >5 MTS. Importantly, calcitriol treatment was able to prevent the increase in the percentage of cells with >5 MTS per metaphase, and the amount of RLF (Figure 3B), indicating a positive role of calcitriol ameliorating genomic instability in VSMC.

Increased genomic instability is frequently associated with cell death. Therefore, to investigate whether RS and genomic instability are promoting VSMC death, we used a trypan blue exclusion assay. Live cells, with intact membranes, exclude trypan blue whereas dead cells do not. The same number of MOVAS were stained using trypan blue every three days for 2 weeks to assess cell viability. MOVAS-GFP or MOVAS-GFP-progerin with different treatments (PDGF-BB or calcitriol) were compared (Figure 3C and 3D). Progerin-expressing cells exhibit decreased viability visualized by an increased number of trypan blue positive cells. On average, the cultures of progerin-expressing cells had three times more dead cells in every count than control cells. In contrast, we did not find an increase in cell death in PDGF-BB treated cells, indicating that cells have mechanisms to cope with the RS induced by the phenotypic switch and prevent genomic instability and the associated cell death. Progerin, in contrast, causes robust RS during the phenotypic switch that hinders proliferation and cellular viability. Interestingly, calcitriol reduces cell death in progerin-expressing cells visualized by trypan blue staining, highlighting the beneficial effect of this hormone in rescuing RS and cellular health (Figure 3D).

Overall, our findings support a detrimental role of progerin in VSMC, promoting RS, fork stalling, and telomere fragility, which compromise cell viability. Interestingly, calcitriol was able to rescue replication defects and reduce genomic instability and cell death in progerin-expressing cells, stressing the potential use of calcitriol as a therapeutic approach to target VSMC loss.

### Progerin induces upregulation of a profibrotic and proinflammatory aging-associated secretory phenotype in AoSMC

To further investigate potential pathways involved in progerin-induced VSMC phenotypic switch and its downstream effects, we examined the transcriptome of two independent experiments of retrovirally transduced human AoSMC (Flag and Flag-progerin), each in duplicate (n=4) using bulk RNA-sequencing analysis (RNA-seq). Gene ontology (GO) analysis of differentially expressed (DE) genes in AoSMC-Flag versus AoSMC-Flag-progerin (fold change (FC) > 1.5 or < -1.5 and p ≤ 0.05), using functional annotation tools DAVID and STRING, revealed strong enrichment of genes in categories such as secretory pathway, extracellular matrix interaction and VSMC differentiation (Figure 4A). This transcriptome analysis confirms that progerin induces a VSMC phenotypic switch, with a significant downregulation of contractile proteins such as α-SMA (ACTA2), myocardin (MYOCD), h1-calponin (CNN1), Myosin-heavy chain (MYH11), smoothelin (SMTN), and caldesmon (CALD1) (Figure 4B). Moreover, most of the DE genes included in ECM interaction and secretory categories are collagens (COLAs), several Cell-ECM adhesion molecules from the integrin superfamily (ITGA/Bs), metalloproteinase(s) (MMPs), and ADAM metallopeptidase Domain(s) (ADAMs) (Figure 4B), indicating a plausible association with abnormal ECM production and organization.

**Figure 4.**
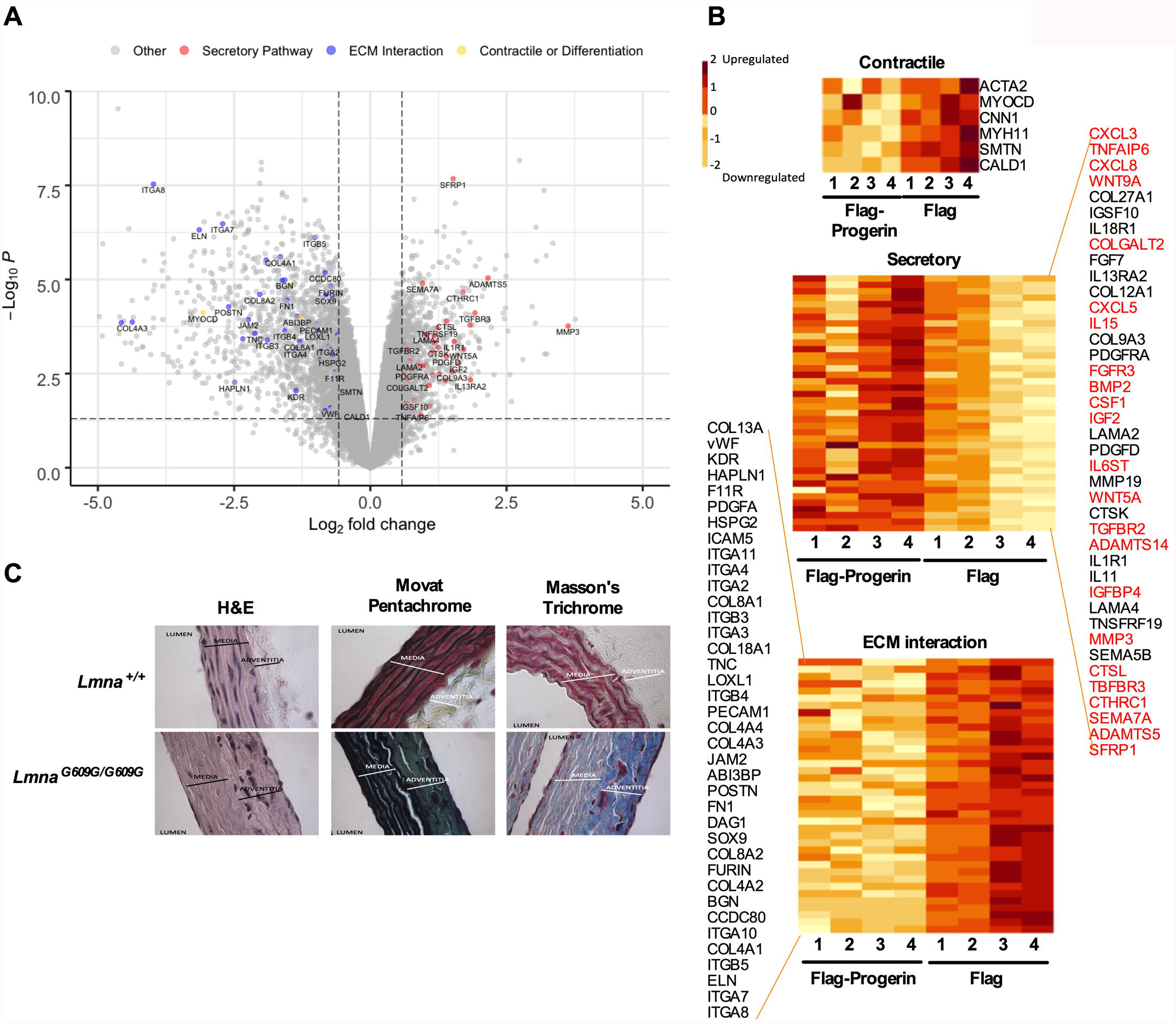
Progerin induces upregulation of profibrotic and proinflammatory aging associated secretory phenotype in AoSMC. A) Volcano plot of Log_2_Fold change (FC>1.5 and < -1.5) (x-axis) and -Log_10_P (y-axis) (FC>1.5 and < -1.5). Note that there are highlighted the main DE genes enriched in the three categories (Secretory, ECM interaction, and Contractile/ Differentiation). B) Heatmap showing clustering of differentially expressed (DE) genes in AoSMC Flag/Flag-progerin. B) Heatmaps showing clustering of differentially expressed (DE) genes in AoSMC Flag/Flag-progerin belonging to three categoryIes: contractile/differentiation (top), secretory pathway (middle), and ECM interaction (bottom). Note that genes associated with the activation of SASP are highlighted in red. C) Representative images from ascending aortas of *Lmna*^*+/+*^ and *Lmna*^*G609G/G609G*^ stained with hematoxylin/eosin (H&E), Movat Pentachrome, and Massons Trichrome’s.

The senescence-associated secretory phenotype (SASP) is characterized by the secretion of prominent levels of inflammatory cytokines, growth factors, and proteases that play a major part in the inflammation associated with age, known as inflammaging. Progerin-expressing endothelial cells were shown to upregulate pro-inflammatory factors associated with SASP (Manakanatas et al., 2022). Using the International Cell Senescence Association Discover Genes Related to Senescence (Senequest) database, we find that the secreted proteins identified in progerin-expressing VSMC belong to the SASP (Figure 4B). These molecules are highlighted in red and correspond to more than 50% of the progerin-induced secreted proteins, indicating that progerin expression in VSMC is able to trigger a senescence-associated secretory phenotype.

ECM abnormalities and secretion of pro-inflammatory factors can result in many pathological processes including tissue fibrosis. We have analyzed the aortas of *Lmna*^*G609G/G609G*^ mice fed a high-fat diet (HFD). These mice live longer than the same mice on regular chow and during their extended lifespan, they develop phenotypes that more closely resemble human HGPS vascular disease. We stained the aortas with Movat Pentachrome and Trichrome staining, well-known methods to evaluate the extent of tissue fibrosis. Results show an evident increase in mucins, collagens, proteoglycans, indicating a fibrotic aorta. Interestingly, there is also a robust loss of VSMC in the medial layer, visualized by a decrease in the number of nuclei in the Hematoxylin-Eosin (H&E) staining (Figure 4C). This massive loss of VSMC in progeria mice has been reported by other investigators (Del Campo et al., 2019; Hamczyk et al., 2018; Sánchez-López et al., 2021). We find that in *Lmna*^*G609G/G609G*^ mice fed a HFD, which live longer, the loss of VSMC, the thickening of the adventitia, and the overall increased fibrosis in the aorta is more dramatic than in progeria mice on a regular diet.

Altogether, these data indicate that progerin expression in AoSMC activates a secretory pathway associated with a profibrotic and pro-inflammatory phenotype, suggesting a key role of proliferative/dedifferentiated VSMC in promoting ECM abnormalities and vascular stiffness in HGPS.

## DISCUSSION

Understanding the mechanisms underlying progerin toxicity in the vasculature is crucial to find therapeutic strategies that ameliorate CVD in HGPS patients. VSMC are extremely sensitive to progerin accumulation, which can affect vessel wall integrity and repair, and play a significant role in CVD in HGPS (Hamczyk et al., 2018; Kim et al., 2021). This work provides new insights into progerin-induced vascular disease. We find that progerin expression in VSMC induces a phenotypic switch towards a synthetic/proliferative phenotype. On one hand, this phenotypic switch triggers gene expression that contributes to ECM dysfunction and vascular stiffness. On the other hand, given the inability of VSMC to properly replicate because of the expression of progerin, forced proliferation of synthetic VSMC has dramatic consequences, including replication stress, telomere dysfunction, increased genomic instability, and cell death.

Several studies have shown that a subset of pathological VSMC-derived cell populations express reduced levels of the contractile VSMC and can undergo a phenotypic switch to mesenchymal-like, fibroblast-like, macrophage-like, osteogenic-like, and adipocyte-like phenotypes that contribute to CVD (Yap et al., 2021). In this work, we show that progerin triggers a phenotypic switch towards a synthetic/fibroblast-like phenotype characterized by the downregulation of contractile markers. Different lines of evidence have also suggested a switch towards a dedifferentiated phenotype in reprogrammed VSMC from HGPS fibroblasts (iSMC). These studies showed less expression of contractile markers such as MYH11, calponin, and α-SMA in iSMC derived from HGPS fibroblasts compared to iSMC derived from non-HGPS fibroblasts. These data indicate an effect of progerin hindering the acquisition of a fully differentiated phenotype (Bersini et al., 2020; Pitrez et al., 2020; von Kleeck et al., 2021). In addition, phenotypic changes in VSMC are often associated with vascular calcification due to their capacity to differentiate to osteogenic-like cells. Indeed, prelamin A, the protein precursor form of the mature lamin A, promotes VSMC calcification and premature aging (Y. Liu et al., 2013). Accordingly, other authors demonstrated that iSMC from HGPS patients upregulate Bone Morphogenetic Protein 2 (BMP2), a key factor involved in the calcification of blood vessels during aging (Petsophonsakul et al., 2019). Here, we demonstrate a robust downregulation of contractile markers (also confirmed by the RNAseq analysis) (Figure 4), with the loss of a fully differentiated phenotype. Importantly, the contractile/fully differentiated phenotype of VSMCs is associated with abundant production of collagen IV (COLA4) and expression of ITGB/A, which in turn is linked to the expression of myoCD or MYH11 (Steffensen et al., 2021; Steffensen & Rasmussen, 2018; Welser et al., 2007). Progerin-expressing cells showed a downregulation of COLA4, ITGB/A, myoCD, and other contractile genes (Figure 4B-C), linking loss of contractibility with ECM dysfunction.

VSMC phenotypic switch is accompanied by an increase in proliferation and the maintenance of a sustained VSMC proliferative capacity is essential for vascular health and repair. Progerin expression is frequently associated with impaired proliferation and DNA damage in different cellular models and also in VSMC. This may have devastating consequences in VSMC that re forced to replicate during the switch (Chojnowski et al., 2020; Hilton et al., 2017; Kang et al., 2021; Kinoshita et al., 2017b; Kreienkamp et al., 2018). A study showed dramatic progerin-induced replication defects which led to DSB accumulation, activation of the DNA damage response, and early replication arrest in HGPS fibroblasts (Hilton et al., 2017). In addition, we have also previously demonstrated that progerin induces replication fork instability (RFI) in RPE (epithelial cells) and human-derived progerin-expressing fibroblasts (Coll-Bonfill et al., 2020; Kreienkamp et al., 2018). Our studies using DNA fiber assays in VSMC demonstrate that progerin increases replication fork speed, induces RS, and promotes fork stalling. We and others previously reported that RAD51, Fanconi anemia proteins, and other factors involved in DNA repair are downregulated in HGPS patient-derived fibroblasts (d’Adda di Fagagna et al., 2003; Takai et al., 2003). These deficiencies might underlie DNA damage accumulation, impacting replication fork progression and promoting fork collapse. More studies are needed to understand the mechanisms underlying progerin-induced replication defects in VSMC and whether deficiencies in DNA repair factors play a major role in RS during the observed VSMC phenotypic switch.

One interesting finding in this study is the effect of PDGF-BB inducing RS. PDGF-BB causes an increase in fork speed and triggers RS. In contrast to progerin-induced RS, PDGF-BB-induced RS is not accompanied by fork stalling, cell death, or telomere fragility. PDGF-BB upregulates DNA-PKcs, a key factor known to increase the capability of VSMC to repair DNA. This suggests a potential role of this factor in preventing PDGF-BB-induced genomic instability and cell death, despite the presence of RS (Kramer et al., 2020). Conversely, the accumulation of unrepaired DNA in progeria cells could have devastating consequences in a proliferative VSMC. In fact, we show that progerin induces telomere fragility and cell death. Different lines of evidence stress the importance to preserve nuclear envelope integrity, and in HGPS this organization is perturbed, causing depletion of the dNTP pools and telomere dysfunction (Kychygina et al., 2021). In addition, deficiencies on telomere capping proteins such as TRF1 induce telomere dysfunction in progeria and other models (Berti et al., 2013; Cao et al., 2011; McCord et al., 2013). It is possible that loss of telomeric proteins also play a role in progerin-induced telomere dysfunction and VSMC loss in HGPS, although future studies are needed to strengthen these conclusions.

Additionally, telomere shortening and dysfunction induce senescence (d’Adda di Fagagna et al., 2003; Takai et al., 2003). Prelamin A accumulation was not observed in young healthy vessels but was prevalent in medial VSMCs from aged individuals, where it colocalized with senescent and degenerate VSMC (Ragnauth et al., 2010). Interestingly, we show activation of a secretory pathway, including aging-related inflammatory and profibrotic factors in progerin expressing AoSMC at the RNA level. There is evidence that inflammatory age-related features can increase the likelihood of developing pathological fibrosis. Indeed, progerin was found in patients with fibrotic cardiopathies and in normal individuals of advanced age (Messner et al., 2018), linking progerin expression and fibrosis in the cardiovascular system. Similarly, HGPS mice show aortas highly rich in collagens, mucins, and ECM matrix components indicating the activation of a profibrotic secretory phenotype that could be associated with vessel stiffness in HGPS.

Lastly, our study identifies a robust effect of calcitriol in rescuing progerin-induced RS, genomic instability, telomere dysfunction, and cell death in VSMC. We postulate that calcitriol slows down replication, as seen by the reduction of tract length in progerin-expressing cells, allowing more time for repair mechanisms to clear the path of the fork of DNA damage. In addition, since calcitriol upregulates several proteins involved in DNA repair, we suggest that calcitriol could also be ameliorating RS and telomere dysfunction upregulating BRCA1, 53BP1 or RAD51 in VSMC as shown in fibroblasts and in other models (Coll-Bonfill et al., 2020; Gonzalez-Suarez et al., 2011; Kreienkamp et al., 2016, 2018). Importantly, these findings suggest that calcitriol might benefit the cardiovascular complications characteristic of patients with HGPS. However, many questions remain to be addressed with respect to the mechanisms underlying calcitriol benefits.

In conclusion, our findings indicate that progerin expression causes a phenotypic switch in VSMC towards a synthetic/proliferative phenotype, which activates a secretory pathway associated with senescence and profibrotic signatures that may trigger vessel stiffness in HGPS arteries. Moreover, VSMC phenotypic switch induces replication defects including fork stalling, RS, and telomere fragility which could be associated with VSMC loss in HGPS.

## EXPERIMENTAL PROCEDURES

### Cell Culture

Primary Aortic Smooth muscle cells from ATCC (PCS-100-012™) (AoSMC) were cultured with SmbM2 (Lonza) with 1x SmGMTM-2 SingleQuotsTM Supplement Pack (CC-4149).

Mouse Vascular Smooth Muscle cells (MOVAS) from ATCC were cultured following the manufacturer’s protocol. Briefly, Dulbecco’s Modified Eagle’s Medium (DMEM) supplemented with 0.2 mg/ml G –418 and 10% fetal bovine serum (FBS) and 1% antibiotics and antimycotics. Both cell lines were maintained at 37C and 5% CO2.

#### Cellular Models

##### Differentiation

AoSMC at 80% of confluence were treated with TGFβ1 10 ng/ml for 24 hours.

##### Dedifferentiation

Synthetic MOVAS were obtained as previously described (Holycross et al., 1992). Briefly, MOVAS were cultured with D-10 media for 24 hours (DMEM/F12, 10%FBS, 1.6 mM Gultamine, and 1% antibiotics and antimycotics). Then, media was changed to ISFM (DMEM/F12, 1.6 mM Glutamine, 0.2 mM L-ascorbic, 5ug/ml transferrin, 6.25 ng/ml Na-Selenate and 1% antibiotics and antimycotics) overnight (ON) and they were stimulated with PDGF-BB 30 ng/ml for 24 hours to induce VSMC phenotypic switch.

##### Apoptosis

50.000 MOVAS were plated in triplicates, and they were stained by trypan blue every 3 days for 2 weeks and counted using a Neubauer Chambre.

### Constructs and Viral Transduction

Retroviral and lentiviral transductions were performed as previously described (Gonzalez-Suarez et al., 2011). Viral envelope and packaging plasmids were gifts from Sheila Stewart (WUSM) were used to generate different cell lines:

- MOVAS EV/progerin: Retroviral particles of pMXIH-EV/progerin (were a gift from Brian Kennedy (Buck Institute)) were used to infect MOVAS and cells infected were then selected with Hygromycin 200μg/ml.
- MOVAS GFP/progerin: retroviral particles containing pBABE-puro-GFP-progerin which was a gift from Tom Misteli (Addgene plasmid # 17663 and PBABE-puro which was a gift from Hartmut Land & Jay Morgenstern & Bob Weinberg (Addgene plasmid # 1764) were used to infect MOVAS and the cells infected were then selected with puromycin 4μg/ml. After selection, cells were sorted using EDTA, to ensure that the cells are in a single cell suspension, and filtering using 1/2-1/3 the diameter to sort them using BD FACSCanto II.
- AoSMC Flag/Flag-progerin: Retroviral particles containing pLPC-Flag and PLPC-Flag-progerin which were s a gift from Gerardo Ferbeyre (Addgene plasmid # 69059, and PLPC-Flag-progerin #69061) were used to infect AoSMC and the cells infected were then selected with puromycin 4μg/ml.
- AoSMC Inducible progerin-expressing cells: Lentiviral particles containing pLenti-CMV-TRE3GNeo-GFP progerin (gift from Tom Misteli (Addgene plasmid # 118710), pLentiCMV-TRE3G-GFP-laminA (gift from Tom Misteli (Addgene plasmid # 118709)), and the constitutive tetracycline repressor A3 mutant expressing pLenti-CMV-rtTA3 (gift from Eric Campau (Addgene plasmid # 26429) to infect AoSMC cells as described in (Kubben et al., 2016). After infection cells were selected using Hygromycin 200μg/ml and Neomycin 500 μg/ml.

### DNA Fiber Assays

Fiber assays were performed as previously described (Berti et al., 2013), with some modifications. Briefly, asynchronous cells were labeled with thymidine analogs: 20 μM iododeoxyuridine (ldU) pfollowed by 200 μM chlorodeoxyuridine (CIdU). The specific labeling scheme and treatments for each experiment are shown in figure legends. Cells were collected by trypsinization, washed, and resuspended in 100 μl of PBS. Then, 2 μl cell suspension was dropped on a polarized slide (Denville Ultraclear) and cell lysis was performed in situ by adding 8 μl lysis buffer (200 mM Tris-HCl pH 7.5; 50 mM EDTA; 0.5% SDS). Stretching of high-molecular-weight DNA was achieved by tilting the slides at 15-45°. The resulting DNA spreads were air-dried and fixed for 5 min in 3:1 Methanol:Acetic acid and refrigerated overnight. For immunostaining, stretched DNA fibers were denatured with 2.5 N HCl for 60 min, washed 3 × 5 min in PBS, then blocked with 5% BSA in PBS for 30 min at 37°C. Rat anti-CldU/BrdU (Abcam, ab6326) (1:100), chicken anti-rat Alexa 488 (Invitrogen, A21470) (1:100), mouse anti-IdU/BrdU (BD Biosciences, 347580) (1:20) and goat anti-mouse IgG1 Alexa 547 (Invitrogen, A21123) (1:100) antibodies were used to reveal CldU- and IdU-labeled tracts, respectively. Labeled tracts were visualized under a Leica SP5X confocal microscope using 63x oil objective lens (NA 1.40, with LAS AF software), and tract lengths were measured using ImageJ. A total of 100-300 fibers were analyzed for each condition in each experiment. All analyses were performed blinding the samples. Statistical analysis of the tract length was performed using GraphPad Prism.

#### Labeling schemes and treatments

To monitor DNA end resection, cells are labeled with thymidine analogs for equal amounts of time: 20 min IdU followed by 20 min CldU or 40 min IdU followed 40 min CldU in PDGF-BB experiments.

### Immunoblotting

Immunoblotting was carried out by lysing cells in UREA buffer (8 M Urea, 40 mM Tris pH 7.5, 1% NP40), for 10 min on ice. Lysates were centrifuged at 14000 rpm for 12 min at 4°C, and the DNA pellet was removed from the sample. 80 μg of total protein was separated by SDS-PAGE on a 4-15% Criterion TGX Gel (Bio-Rad) and transferred to a nitrocellulose membrane using the Trans-Blot Turbo system (Bio-Rad). Membranes were blocked using 5% BSA in TBS+0.1% Tween-20 for 1 hour at room temperature, then incubated overnight at 4°C with the appropriate antibody diluted in blocking solution. Membranes were washed 3X using TBS+0.1% Tween-20 after both primary and secondary antibody incubations. Membranes were developed using Immobilon Western Chemiluminescent HRP Substrate (Merck Millipore). Antibodies used: Calponin (ab46794) 1/1000, α-SMA (CBL171-I Millipore) 1/1000, and lamin A (ab26300).

### Immunofluorescence

3 × 105 cells were fixed in 3.7% formaldehyde + 0.2% Triton-X100 in PBS for 10 min, washed 3× in PBS, and blocked 1 h at 37 °C in 10% BSA/PBS. Incubations with antibodies were performed ON 4°C, in a humid chamber. After washes in PBS, cells were counterstained with DAPI in Vectashield. Microscopy and photo capture was performed on a Leica DM5000 B microscope using 40× or 63× oil objective lenses with a Leica DFC350FX digital camera and the Leica Application Suite. <u>Antibodies used</u>: Calponin (ab46794) 1/50, α-SMA (Millipore CBL171-I) 1/500, and Ki67 (ab92742) 1/1000.

### Quantitative Reverse-Transcription PCR

cDNA was generated by reverse transcription of 1 μg total RNA using GeneAmp RNA PCR kit. qRT-PCR was performed using the 7500HT Fast Real-Time PCR system with the Taqman Universal PCR Master Mix or Universal SYBR Green Supermix. Reactions were carried out in triplicate and target gene and endogenous controls amplified in the same plate. Relative quantitative measurements of target genes were determined by comparing cycle thresholds.

Primers used: Calponin F CACGACATTTTTGAGGCCAA, Calponin R TTTCCTTTCGTCTTCGCCAT, sm22v2 F GGAAGCCTTCTTTCCCCAGA, sm22v2 R TCCA GCTCCTCGTCATACTTCTT, myoCD F GCACCAAGCTCAGCTTAAGGA, myoCD R TGGGAGTGGGCCTGGTTT. Taqman probe: gapdh Hs02758991 and progerin HsAihsp3z.

### Metaphase spreads

Cells were cultured in 10 mm dishes to 70% confluence and asynchronous cells were exposed to DMSO, 4 mM HU, 50 μM Mirin, or 4 mM HU + 50 μM Mirin for 5 hours. Cells were allowed to complete S-phase in complete medium supplemented with 10 μM Colcemid (Sigma #D1925) in order to arrest them in metaphase. After collecting the culture media, cells were washed with 1X PBS (which was also collected), and the trypsinized cells were collected. All the fractions were combined and centrifuged to pellet the cells. Supernatant was then aspirated to leave ∼2 ml of media + cell pellet, which was resuspended by gentle flicking. 9 ml of pre-heated (37°C) hypotonic buffer (0.56% KCl) was added then to the cell suspension, which was incubated in a 37°C water bath to allow hypotonic swelling of the cells. A small amount (∼500 μl) of fixing solution (3:1 methanol:acetic acid) was added and the cells were pelleted at 4°C by centrifugation. Cells were kept on ice from this point onwards. After aspirating the supernatant to leave ∼2 ml, the cells were resuspended by gentle flicking and 9 ml fixing solution was added dropwise. The suspension was centrifuged once more to pellet the cells and fixing solution added in an equivalent manner. This mixture was stored at -20°C overnight. Cells were then pelleted, resuspended in ∼500 μl of fresh fixing solution, dropped onto microscope slides, and allowed to air-dry at room temperature ON.

### Fluorescence in situ hybridization

FISH on metaphase spreads was performed as described in reference (Samper et al., 2000). In brief, cells were arrested in mitosis by treating with colcemid for 4 h and prepared for FISH by hypotonic swelling in 0.56% KCl, followed by fixation in 3:1 methanol:acetic acid. Cell suspensions were dropped onto slides and FISH was performed using a Cy3-telomeric PNA probe and DNA counterstained using DAPI. Images were visualized under a Leica DM5000 B microscope using 100X oil objective lens (NA 1.3) with a Leica DFC350FX digital camera and the Leica Application Suite (Version 4.1.0).

### Histology

The proximal thoracic aorta was excised, cleaned from surrounding fat tissue, and divided into three fragments: ascending (from heart to brachiocephalic braching), arch (from brachiocephalic to left subclavian braching), and descending (after left subclavian braching). These three segments were fixed 5 days in buffer containing 10% formalin, 5% sucrose, and 2 mmol/L EDTA, and subsequently embedded in paraffin blocks. Four-micrometer sections were then stained with hematoxylin and eosin, following standard techniques to analyze fibrosis.

### RNA Sequencing and Analysis

Samples were prepared according to the library kit manufacturer’s protocol, indexed, pooled, and sequenced on an Illumina NovoSeq 6000. Basecalls and demultiplexing were performed with Illumina’s bcl2fastq software and a custom python demultiplexing program with a maximum of one mismatch in the indexing read. RNA-seq reads were then aligned to the Ensembl release 76 primary assembly with STAR version 2.5.1a1. Gene counts were derived from the number of uniquely aligned unambiguous reads by Subread:featureCount version 1.4.6-p52. Isoform expression of known Ensembl transcripts was estimated with Salmon version 0.8.23. Sequencing performance was assessed for the total number of aligned reads, the total number of uniquely aligned reads, and features detected. The ribosomal fraction, known junction saturation, and read distribution over known gene models were quantified with RSeQC version 2.6.24.

All gene counts were then imported into the R/Bioconductor package EdgeR5 and TMM normalization size factors were calculated to adjust for samples for differences in library size. Ribosomal genes and genes not expressed in the smallest group size minus one samples greater than one count-per-million were excluded from further analysis. The TMM size factors and the matrix of counts were then imported into the R/Bioconductor package Limma6. Weighted likelihoods based on the observed mean-variance relationship of every gene and sample were then calculated for all samples with the voomWithQualityWeights7.

### Study approval

All animal studies were approved (protocol #2299) and conducted in accordance with the Animal Studies Committee at Saint Louis University (Dr. John C. Chrivia, Ph.D. is the Institutional Animal Care and Use Committee Chair)

### Statistical analysis

All data sets from each DNA fiber assay experiment have been first subjected to normality test. Most of the data set does not meet the criteria of a normal distribution. Since each data set includes more than 100 values to calculate the statistical significance of the observed differences we have used: non-parametric tests, such as Wilcoxon matched-pairs signed-rank test for paired samples and Mann-Whitney test for unpaired samples; one-way ANOVA was used for all the experiments in which more than two comparisons were performed. GraphPad Prism 6 was used for the calculations. In all cases, differences were considered statistically significant when p<0.05.

## ACKNOWLEDGEMENT & AUTHOR CONTRIBUTIONS

We thank GTACC Genomic Core from WUSM, and especially Elliot Klotz and Eric Tycksen, for the RNA seq bioinformatic analysis. We acknowledge Michelle Pherson from SLU Genomic Core for providing help with RNA seq analysis, generating Heatmaps and Volcano plots. N.C.B. performed the design, most of the experiments, and wrote the manuscript. U.M. performed some of the experiments. C.J.L. and R.P.M. performed all the analysis of aortic samples and provided insightful comments. S.G. supervised the research and the writing of the manuscript.

## NO CONFLICT OF INTEREST

